# Sex-specific bone marrow populations regulate skeletal phenotype and identify CD74-marked osteoclast precursor subsets

**DOI:** 10.64898/2026.11.03.763975

**Authors:** Jinsha Koroth, Ismael Y. Karkache, Elizabeth K. Vu, Joseph C. Manser, Samuel Mitchel, Mitchell J. Shimak, Nole Robertson, Emily Patterson, Sydney Kiraguri, Kim C. Mansky, Elizabeth W. Bradley

## Abstract

While there are clear differences in bone accrual and maintenance between men and women due to hormonal regulation, the potential cell-intrinsic factors governing this difference are poorly understood. To investigate this, we performed bone marrow transplantation assays using male or female bone marrow transplantation into male hosts. Micro-CT analyses of lethally irradiated 8-week-old male mice transplanted with female bone marrow exhibited increased vertebral body trabecular and cortical bone mass. To identify cell types mediating these effects, we performed comparative single cell sequencing analyses and identified several CD11B^+^CD74^+^ myeloid cell clusters. Male donor BM chimeric mice had a higher proportion of Cd74-expressing clusters compared to female donor BM chimeric mice. We confirmed that males have greater numbers of bone marrow-resident CD11B^+^CD74^+^ cells under physiological conditions. Elevated expression of X-linked genes *Tlr7* and *KdmCa* marked a population of CD11B^+^CD74^+^ cells. Lastly, we show that female CD11B^+^CD74^+^ cells exhibit diminished osteoclastogenesis. These findings demonstrate that male and female bone marrow-resident cells have cell-intrinsic differences that regulate bone mass and that CD74 marks a subset of osteoclast precursor–enriched myeloid populations.

## Introduction

Bone accrual and maintenance differ between men and women. Women achieve peak bone mass at a younger age than men, but men exhibit greater bone mass over their lifetime. Bone loss in both men and women starts prior to sex-steroid insufficiency (e.g., menopause), but women experience a larger percent loss during their midlife years (1). The structural basis for bone loss during midlife likewise differs between men and women, with bone loss in women characterized by reductions in trabecular number and bone loss in men due to diminished trabecular thickness (2). Women also experience precipitous bone loss following the menopause that can be mitigated by estrogen replacement therapy (3). Murine models likewise recapitulate these findings (4). Divergences in bone accrual and maintenance between men and women translates to an increased risk of osteoporosis and fragility fracture amongst women; conversely, mortality is greater for men following a fragility fracture.

Circulating sex hormones are well established as regulators of bone mass in both men and women. Estradiol (E2) limits osteoclast-mediated bone resorption and declining estrogen levels lead to enhanced bone loss in both humans and pre-clinical models (3). Men are not estrogen deficient, but have comparably lower levels of circulating estrogens (3). Estrogens are crucial for maintaining bone health in men as well as women (3). Importantly, female cells have increased estrogen receptor (α/β) density and respond with greater amplitude to E2 (5). Targeted deletion of ER-α/β in osteoclast-lineage cells decreases cancellous bone mass (6), but local production of estradiol by bone cells (i.e., via aromatase) may also influence bone remodeling (7).

While circulating sex hormones regulate bone remodeling, cell-intrinsic factors may likewise dictate divergences in bone accrual and maintenance between men and women. Prior reports demonstrate that male and female cells exhibit cell-intrinsic epigenetic differences that are independent of circulating sex hormones (8). Two publications using the four core genotypes model, in which chromosomal sex can be separated from gonadal sex, also demonstrate that cell-intrinsic factors may facilitate alterations in bone remodeling between males and females (9, 10). In mice harboring ovaries, bone mass was enhanced with an XX chromosomal content as compared to those with XY (10), demonstrating that cell-intrinsic differences due to chromosomal content control bone mass. Despite these advances, our understanding of the cellular and molecular mechanisms that govern cell-intrinsic regulation of bone remodeling is poorly defined.

The overall goal of our study is to define how cell-intrinsic factors of bone marrow-resident hematopoietic lineage cells affect bone remodeling and vertebral body bone mass. We used the model of transplantation of female bone marrow into male hosts coupled with single cell sequencing as a model to uncover potential cell-intrinsic factors governing bone remodeling.

## Methods

### Animal Procurement and Housing

The Jackson Laboratory provided male and female CD45.2^+^ C57Bl/6J and CD45.1^+^ B6.SJL-Ptprca Pepcb/BoyJ mice (strain #000664 and #002014, respectively, Bar Harbor, ME) aged to either 4-6-weeks old. Mice were then acclimated for two weeks prior to study initiation. All animal research was conducted according to guidelines provided by the NIH and the Institute of Laboratory Animal Resources, National Research Council. The University of Minnesota IACUC approved all animal studies.

### Whole Bone Marrow T Cell Depletion

Hind limbs from male and female 6-8-week-old C57BL/6J mice were euthanized by CO_2_ asphyxiation and bone marrow was flushed with sterile PBS. Bone marrow cell suspensions were pooled, pelleted, and the red blood cells were lysed (1X RBC lysis buffer, #00-4333-57, ThermoFisher Scientific, Waltham, MA). Nucleated cells were pelleted and suspended in MACS buffer. PE-conjugated anti-CD8 and anti-CD3 antibodies (Miltenyi Biotech, #130-102-807 and #130-121-133) were added to label T cells. Cells were then incubated with anti-PE microbeads (Miltenyi Biotec, #130-048-801), washed, and then applied to an LS column (Miltenyi Biotec, #130-042-401) placed in a MidiMACS Separator magnet. Eluted CD8^-^/CD3^-^ bone marrow cells were collected in PBS and used for bone marrow transplantation assays.

### Irradiation and Bone Marrow Transplantation

Six to 8-week-old male mice (n=18) were lethally irradiated using two 500 rad doses separated by a period of 4 hours and then transplanted with T cell-depleted bone marrow derived from either age-matched males (n=10) or age-matched females (n=8). Following bone marrow transplantation, mice were housed in autoclaved cages and administered polymyxin B and neomycin sulfate for two weeks. Mice were aged to 24-weeks-old and then euthanized for analyses. All animals were included within the study.

### Micro-Computed Tomography

Femora and spines were collected from 24-week-old chimeric mice and fixed in 10% neutral buffered formalin for 48 hours then stored in 70% ethanol. Blinded scanning was performed using the XT H 225 micro-computed tomography (micro-CT) machine (Nikon Metrology Inc., Brighton, MI) set to 120kV, 61µA, 720 projections at two frames per projection with an integration time of 708 milliseconds. Scans were done at an isometric voxel size of 7.11 µm with a 1mm aluminum filter. Each scan volume was reconstructed using CT Pro 3D (Nikon metrology, Inc., Brighton, MI) and then converted to bitmap datasets using VGStudio MAX 3.2 (Volume Graphics GmbH, Heidelberg, Germany). Scans were reoriented via DataViewer (SkyScan, Bruker micro-CT, Belgium) to create a new bitmap dataset. Morphometric analysis was performed using SkyScan CT-Analyzer (CTAn, Bruker micro-CT). 3D Analysis of trabecular bone was performed in the distal femoral metaphysis 0.7 mm proximal to the growth plate and extending 1.5 mm proximally toward the bone diaphysis. For the femoral cortex, 2D analysis occurred in a 0.5 mm section within the mid-diaphysis defined as 4 mm from the growth plate. Regions of interest were set by automated contouring for the trabecular and cortical ranges, with some manual editing when necessary. For the spines, 3D analysis was conducted in the vertebral body of the L4 vertebrae of the mouse spine. The inner trabecular bone was delineated from the surrounding vertebral processes by consistent manual assignment of an elliptical ROI, formed between the superior vertebral endplate extending to the beginning of the inferior endplate, with a new ROI manually generated every 10 images within a dataset. CtAn interpolated the ROI for the remaining images. Vertebral cortical bone was delineated by inverting the trabecular ROI of the isolated vertebral bodies. For each sample, 3D analysis of the trabecular and cortical bone was performed at a globalized threshold and Cortical and Trabecular Thickness (Ct.Th/Tb.Th), BV/TV (Ct.BV/TV/Tb.BV/TV), Trabecular Separation (Tb.Sp), and Trabecular Number (Tb.N) were measured. 3D models of the ROIs were reconstructed using CT Vox for qualitative analysis of bone.

### Bone Histomorphometry

Following micro-CT analyses, vertebrae from chimeric mice (n=5/group) were decalcified in 15% EDTA for 14 days. Tissues were paraffin embedded and 7-μm sections were collected and stained with TRAP/Fast Green (Sigma, number 387A-1KT) or hematoxylin/eosin. Standardized histomorphometry to quantify bone volume per total volume (BV/TV) as well as the number of osteoclasts and osteoblasts per bone perimeter (No. Oc/B.PM, No.Ob/B.PM, respectively) was performed (11).

### Magnetic Assisted Cell Sorting and Flow Cytometry

Bone marrow cells were flushed from the hind limbs of 24-week-old chimeric mice. Red blood cells were lysed, cells were pelleted, enumerated, and then suspended in MACS buffer. CD11B^+^ cells were then collected by magnetic assisted cell sorting as previously described (12). For isolation of CD74^+^ cells, cells were then incubated with Allophycocyanin (APC)-conjugated anti-CD74 (RCD Systems, #FAB7478A) antibodies. Cells were pelleted, washed, suspended in 1X MACS buffer and incubated with anti-APC microbeads (RCD Systems, #FAB7478A). Cells were washed and then isolated are previously described (12) using anti-APC beads (Miltenyi Biotec, #130-090-855). For assessment of chimerism, we prepared single stained controls and double stained samples (*n* = 3) with unsorted nucleated cells with BV421-conjugated anti-CD45.1 (Biolegend, #110731), APC-conjugated anti-CD45.2 (Miltenyi Biotech, #130–102-964), or co-stained with anti-CD45.1 and anti-CD45.2 antibodies, respectively. Data were acquired on a Fortessa X-20 and analyses were performed using BD FACSDiva and FlowJo software.

### Single Cell RNA-Sequencing

CD11B^+^ cells from chimeric mice in each group (n=3) were isolated via MACS as described above and pooled. Single cell cDNA libraries from each pooled sample were then prepared using Chromium Single cell 3′ Reagent v3 Kits (10× Genomics, Pleasanton, CA, USA) according to the manufacturer’s protocol. Quality of amplified cDNA was assessed using an Agilent Bioanalyzer. Following quality control, the cDNA libraries were sequenced on an Illumina NovaSeq 6000 sequencer at a depth of 50,000 reads per cell. The Genomics Core at the University of Minnesota performed library construction and sequencing.

### Single Cell Bioinformatic Analyses

10× Genomics Cell Ranger software (version 6.0.0) was used to process the raw scRNA-seq data in accordance with the guidelines specified by the manufacturer. Cell Ranger software (2020-A, 10× Genomics Cell Ranger Count v7.0.1) was used to demultiplex the raw base call (BCL) files produced by the Illumina NovaSeq 6000 sequencer. Next, Cell Ranger was used to count the unique molecular identifiers (UMI) and barcodes and align fastq files to the mouse reference genome (GRCm39). Quality assurance and subsequent studies were then performed on the feature-barcode matrices created by Cell Ranger. The study was filtered to exclude cells with outlier status, aberrant gene detection rates (<500 and >5,000), and high mitochondrial transcript level (>8%, a sign of cellular stress). Uniform manifold approximation and projections (UMAPs) were used to generate non-linear dimensional reductions and identification cell clusters. Cell Ranger and Cloupe Browser software was used for data integration comparing between samples (10x Genomics).

### Bioinformatic Analyses

Resulting fold changes from scRNA-Seq analyses were filtered for differentially expressed genes with a p<0.05 and counts greater than zero and used in tertiary bioinformatic analyses. Functional annotation data were generated using the Gene Ontology Resource GO Biological Processes SLIM (GO, geneontology.org). Heatmaps of resulting fold changes were generated using Morpheus (software.broadinstitute.org/Morpheus).

### Osteoclastogenesis

Osteoclasts were generated as previously described (13). Briefly, the hind limbs of 6–9-week male or female mice were collected and the bone marrow was flushed. Red blood cells were lysed in RBC lysis buffer and nucleated cells were MACS sorted for CD11B^+^CD74^+^ and CD11B^+^CD74^-^ populations as described above. Following MACS, 0.4×10^6^ cells/cm^2^ from each sorted population were cultured in phenol-red-free alpha minimal essential medium supplemented with 10% heat-inactivated FBS, 100 ng/ml RANKL (#315-11-50UG, Peprotech) and 50 ng/ml M-CSF (#416-ML-050, RCD Systems). Medium and cytokines were refreshed on day 3 and cells were fixed and TRAP stained (#387A-1KT, Sigma Aldrich) on day 4.

### Semi-Quantitative PCR

Total RNA was extracted using TRIzol and chloroform. Reverse transcription using the iScript™ Reverse Transcription Supermix (Biorad, #1708841) using 1 μg of total RNA was used to generate cDNAs. Gene expression was then assessed via real-time PCR using gene-specific primers listed in **Supplementary Table 1**. Fold changes in gene expression for each sample were calculated using the 2^−ΔΔCt^ method relative to control after normalization of gene-specific C_t_ values to Ywhaz C_t_ values (14, 15). Shown are averaged data from three independent replicate experiments.

### Flow Cytometry

Bone marrow cells were isolated as previously described. For flow cytometry analysis of CD11B^+^CD74^+^ cells, we prepared single stained controls and double stained samples (n=3) with unsorted nucleated cells with PE-conjugated anti-CD11B (Miltenyi Biotech, #130-113-235), APC-conjugated anti-CD74 (RCD Systems, #FAB7478A), or co-stained with anti-CD11B and anti-CD74 antibodies, respectively. Data were acquired on a Fortessa X-20 and analyses were performed using BD FACSDiva^TM^ and Flowjo software.

### Statistics

Statistics were performed in GraphPad Prism (Version 10) using Student’s t-test or one-way ANOVA as appropriate and post-hoc tests for multiple comparisons when necessary. Specific *p* values are shown within figures, with *p*<0.05 considered as significant. Data are shown as box plots from the 25th to 75th percentiles, with whiskers extending to the minimum and maximum value and means shown by horizontal lines or means ± standard deviation.

## Results

To understand how cell-intrinsic factors may dictate differences in bone remodeling between men and women, we lethally irradiated 6-8-week-old male mice and then transplanted T-cell depleted bone marrow cells derived from age-matched male or female mice (Figure 1A). Recipient mice were then aged to 24-weeks. We next evaluated percent chimerism within each group. Within CD45.1 hosts, 90% of bone marrow cells were CD45.2^+^, indicating nearly complete chimerism. (Figure 1B). We did not observe a difference in body weight or spleen weight to body weight between groups (Figure 1C, D).

**Figure 1.**
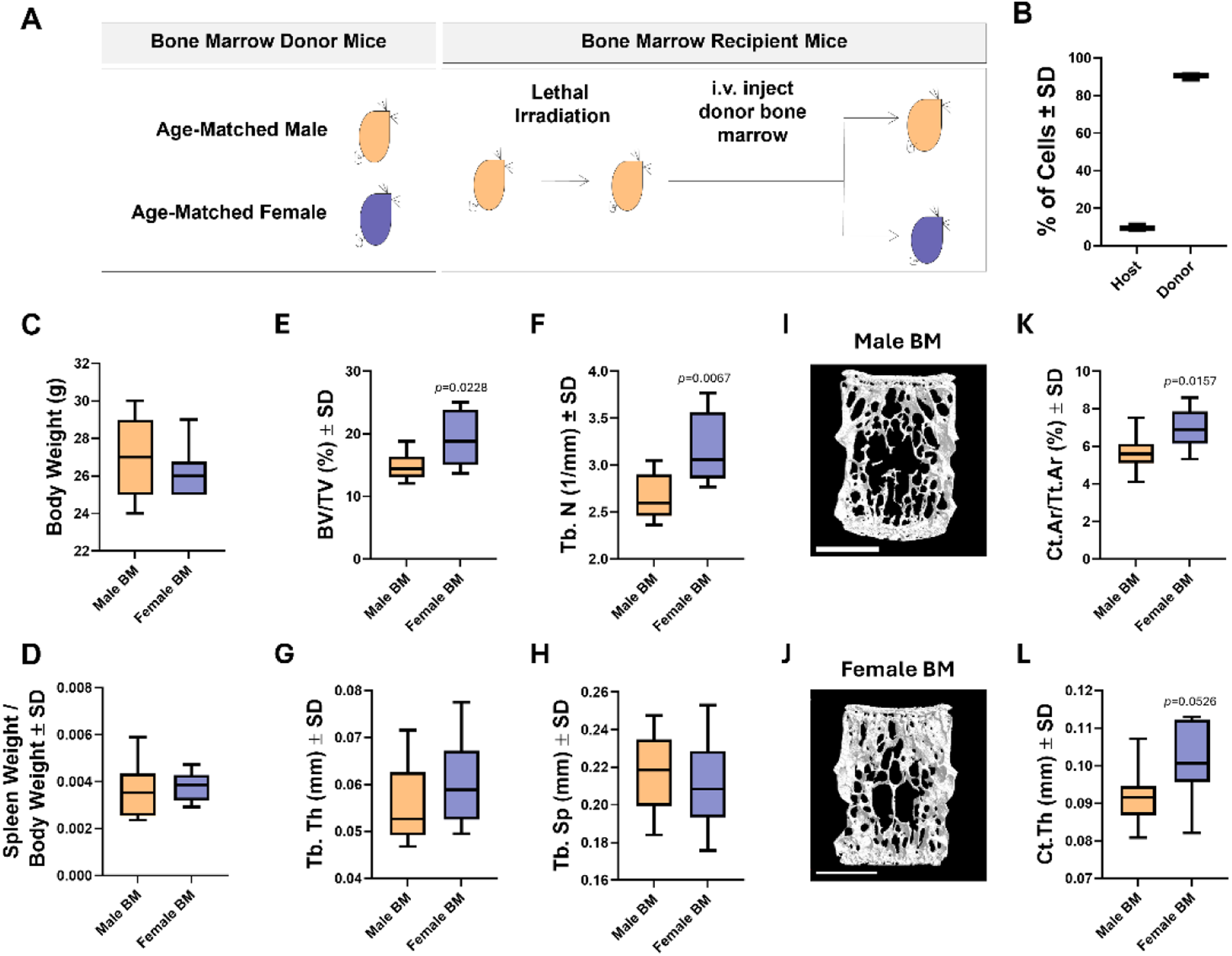
Transplantation of female cells into male hosts enhances vertebral body bone mass. (A) Overview of experimental design. Male hosts were lethally irradiated, transplanted with either T cell-depleted male or female bone marrow, and aged to 24-weeks-old. (B) Percent chimerism. (C) Body weight. (D) Spleen weight to body weight ratio. (E-L) Micro-CT analysis of the fourth lumbar vertebral body was performed. (E) Bone volume per total volume (BV/TV), (F) Trabecular number (Tb. N), (G) Trabecular thickness (Tb. Th), (H) Trabecular spacing (Tb. Sp), (I-J) reconstructions, (K) Cortical area per total area (Ct.Ar/ Tt.Ar) and (L) cortical thickness (Ct. Th) are shown. *p* values under 0.05 are reported.

We next assessed changes to vertebral bone mass. Micro-computed tomography scanning revealed that male hosts transplanted with female cells exhibited a 30 percent increase in vertebral bone volume per total volume (Figure 1E, I, J). This was accompanied by an increase in trabecular number (Figure 1F), but no changes to trabecular thickness or spacing were observed (Figure 1G, H). Vertebral body cortical bone mass and cortical thickness were also elevated in mice transplanted with female bone marrow (Figure 1K, L). T cells, including CD4^-^CD3^+^ cells, express RANKL (16). Prior work demonstrates that male and female C57Bl6 mice do not demonstrate altered numbers of CD3^+^/CD8^+^ T cells when comparing males and females (17), which we confirmed in our analyses (Supplemental Figure 1A). Male and female CD4^-^CD3^+^ T cells could also exhibit differential expression of RANKL. Isolated CD3^+^CD8^+^ T cells from male and female mice did not demonstrate differential expression levels *Tnfsf11* / RANKL levels (Supplemental Figure 1B). We confirmed enhanced vertebral body BV/TV via standardized bone histomorphometry (Figure 2). While there was a trend towards diminished osteoclasts per bone perimeter in mice transplanted with female bone marrow, this effect was not significant (Figure 2B). The number of osteoblasts per bone perimeter was likewise not altered (Figure 2C).

**Figure 2.**
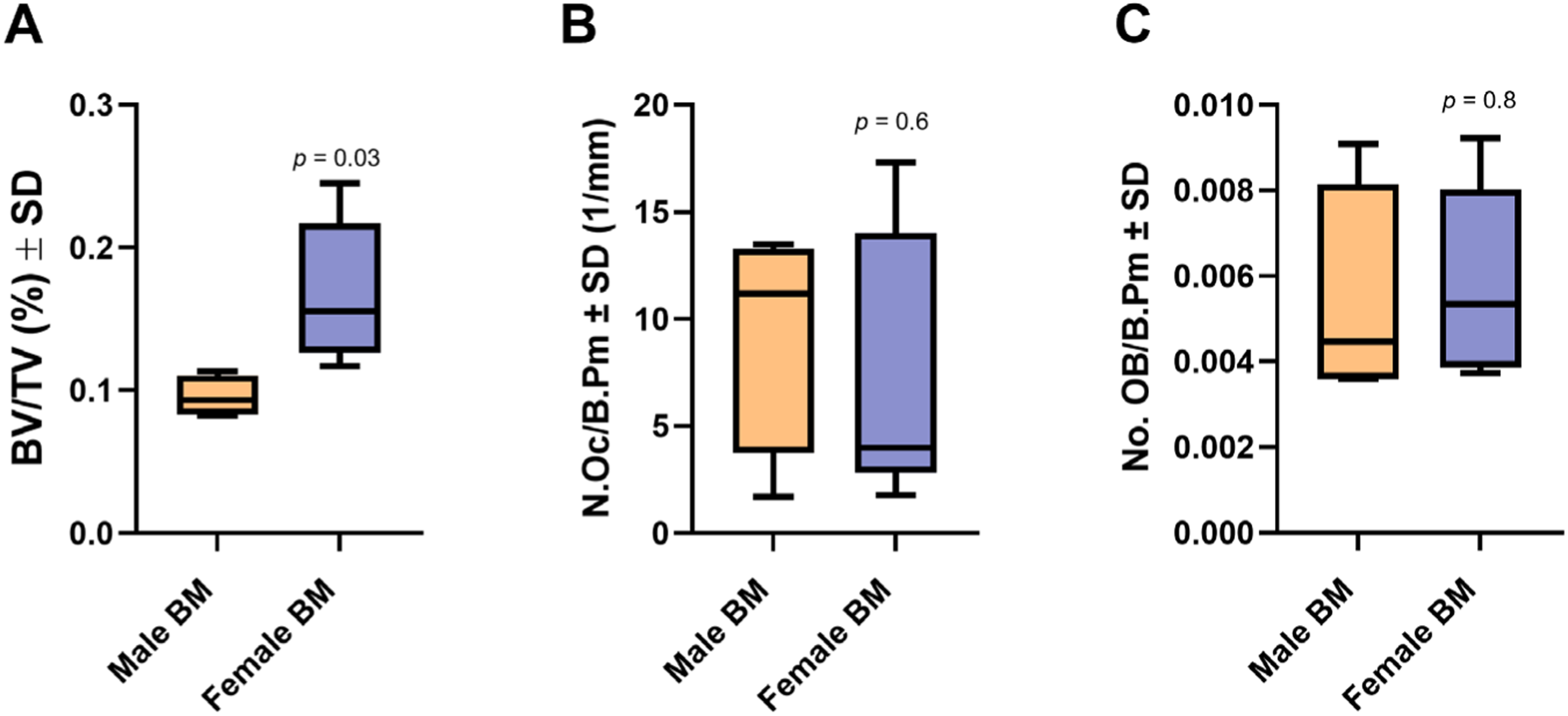
Bone Histomorphometry following transplantation of female cells into male hosts. Male hosts were lethally irradiated, transplanted with either T cell-depleted male or female bone marrow, and aged to 24-weeks-old. Spines were decalcified, embedded, and sectioned. TRAP/Fast green staining and HCE staining was performed. The (A) BV/TV, (B) the number of osteoclasts per bone perimeter and (C) the number of osteoblasts per bone perimeter within the L4 vertebral body was evaluated by histomorphometry.

Given the changes in vertebral body bone mass, we also assessed bone mass within the appendicular skeleton. Micro-CT scanning of the distal femur did not reveal a change in distal femoral trabecular bone volume per total volume (Supplemental Figure 2A, D). There was no change in femoral trabecular thickness, but trabecular number increased with a corresponding decrease in trabecular spacing (Supplemental Figure 2B-E).

To identify potential mechanisms governing the increase in bone mass, we performed single-cell RNA sequencing (scRNA-Seq) of CD11B^+^ bone marrow cells isolated from chimeric mice. UMAP representations of the scRNA-Seq data revealed 16 distinct clusters (Figure 3A). Although all clusters were represented in both samples, the cellular distribution within clusters differed. Cell numbers within clusters 2, 8, 12, 13, 15, and 16 were enriched within male hosts transplanted with male cells. Clusters 2 and 13 did not exhibit any significant differential gene expression, but contained Ly6g^+^Csf3r^+^ neutrophils and Ly6c^+^Cxcl3^+^ monocytes respectively. Cluster 8 was defined by expression of *LyCg* and *LyCc1* and may represent immature neutrophils (Figure 3A, B). Cluster 12 was defined by expression of *Cd34*, *Il2rb*, and *Elane*, potentially representing promyelocytes (Figure 3A, B). Clusters 13 and 15 were characterized by expression of *LyCc1*/*Cxcl3* and *Cd74*/*Cd20Sa*/*Itgax*/*Ccr5*, respectively (Figure 3A, B). We also noted expression of osteoclast precursor genes, including *Tnfrsf11a*/RANK, *Acp5*/TRAP, *Cx3cr1*, by cells within clusters 11 and 14 that were enriched in male hosts transplanted with female bone marrow (Figure 3A, B). Clusters 11, 14, 15, and 16 all showed enrichment in *Cd74* expression (Figure 3A, B). Hierarchical clustering based on phenotypic markers also demonstrated high similarity between clusters 11, 14, 15, and 16 (Figure 3B). *Cd74* was the most variable feature (Figure 3C, D). The top five upregulated and downregulated genes within each cluster across all clusters are shown in Figure 3E and F, respectively.

**Figure 3.**
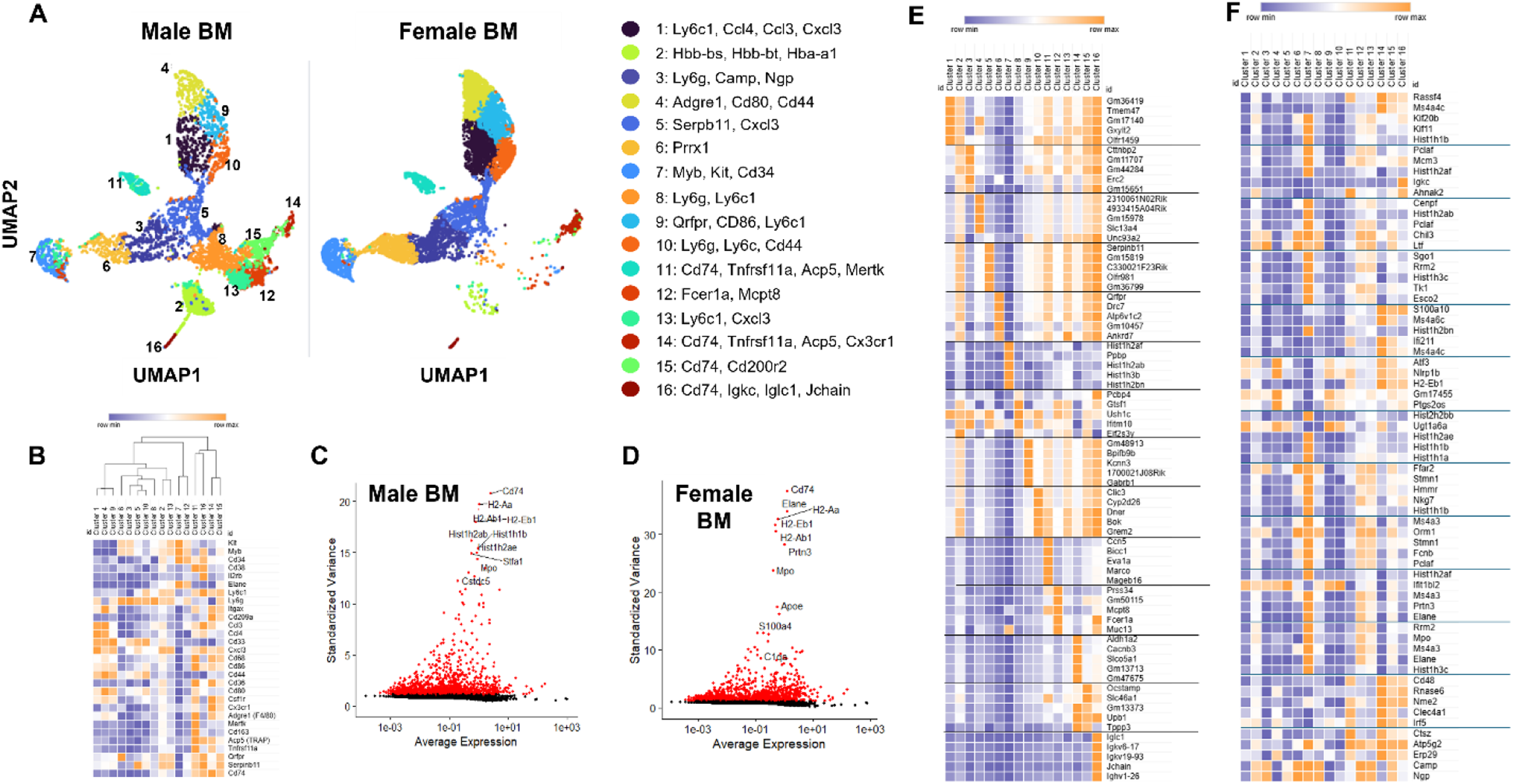
Single-cell RNA-Sequencing of CD11B+ bone marrow cells from chimeric mice. (A-D) Male recipient mice were lethally irradiated transplanted with T cell-depleted bone marrow (BM) from age-matched males or age-matched female donor mice. Recipient mice were aged to 24-weeks for subsequent analyses. CD11B^+^ cells were collected from mice and scRNA-Seq was performed. Resulting data were integrated and comparative analyses were performed. (A) UMAP 2D representation of CD11B^+^ clusters derived from recipient mice. (B) Heatmap of phenotypic genes used for characterization. Significantly variable features from male (C) and female (D) scRNA-Seq datasets. (E) Heatmap of the top five upregulated genes within each cluster across all clusters. (F) Heatmap of the top five downregulated genes within each cluster across all clusters.

Because of the differential cellular distribution and potential to regulate the bone remodeling process, we performed subsequent bioinformatic analyses on clusters 11 and 14. The top 20 most differentially regulated genes within clusters 11 and 14 across all clusters are illustrated in Figures 4A and 5A, respectively. Likewise, surface markers defining these clusters are shown in the heatmaps in Figures 4B and 5B. GO analysis of differentially expressed genes by cells within cluster 11 was also performed. Fold enrichment for functional annotation categories and bubble plots representing these analyses for cluster 11 are shown in Figure 4C, D. Corresponding data for cluster 14 are depicted in Figure 5 C, D. GO analysis of cluster 14 demonstrated enrichment for genes associated with immunity, but also for regulation of osteoclast differentiation (Figure 5C, D).

**Figure 4.**
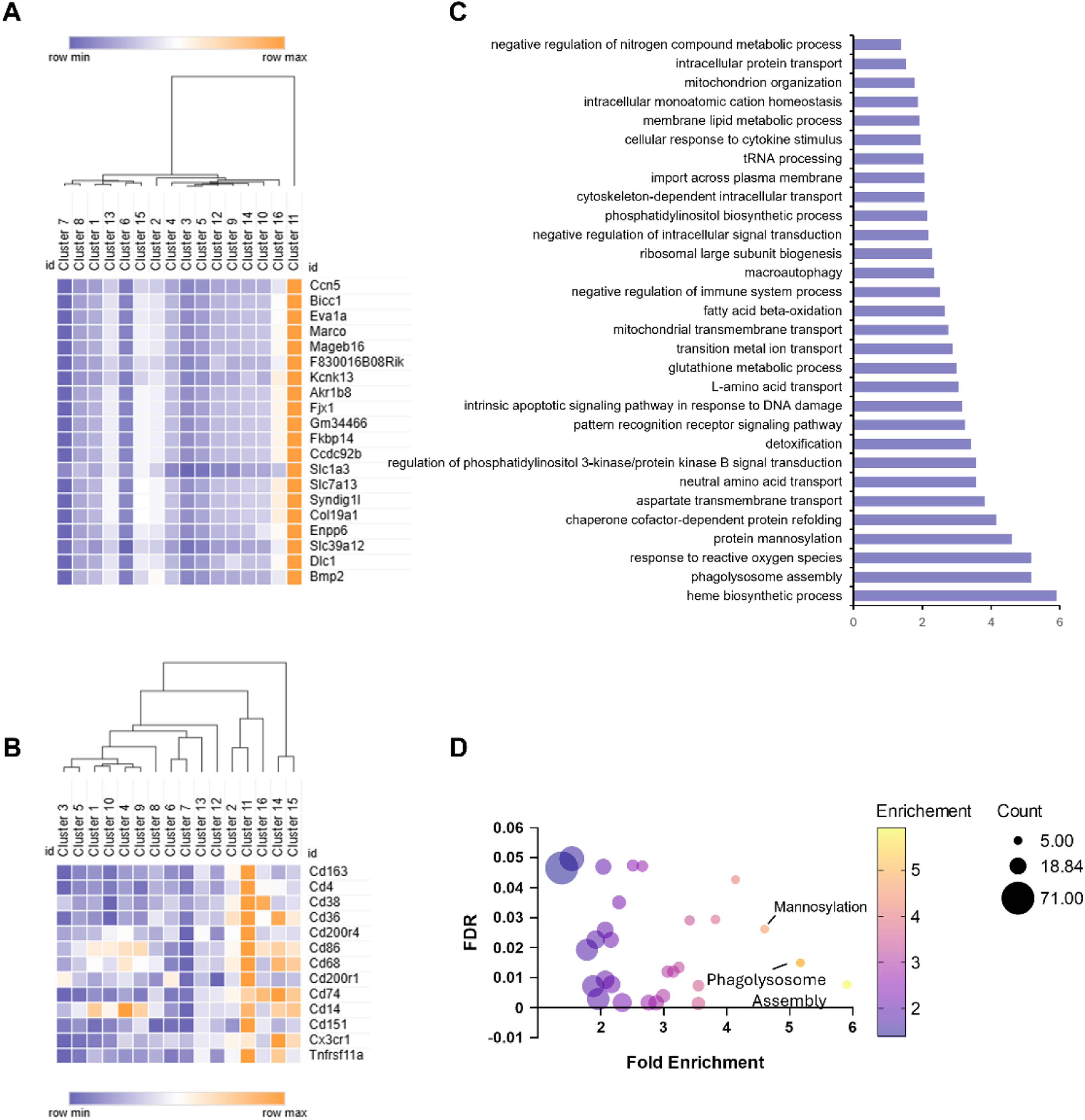
Cluster 11 Bioinformatic Analysis. (A) Heatmap of the top 20 differentially expressed genes within cluster 11 across all clusters. (B) Heatmap of surface markers defining cluster 11 across all clusters. (C) GO biological process SLIM analyses were performed using differentially expressed genes within cluster 11. Shown are enrichment scores for each functional annotation. (D) Bubble plot of enrichment, gene counts, and the false discovery rate for each functional annotation term.

**Figure 5.**
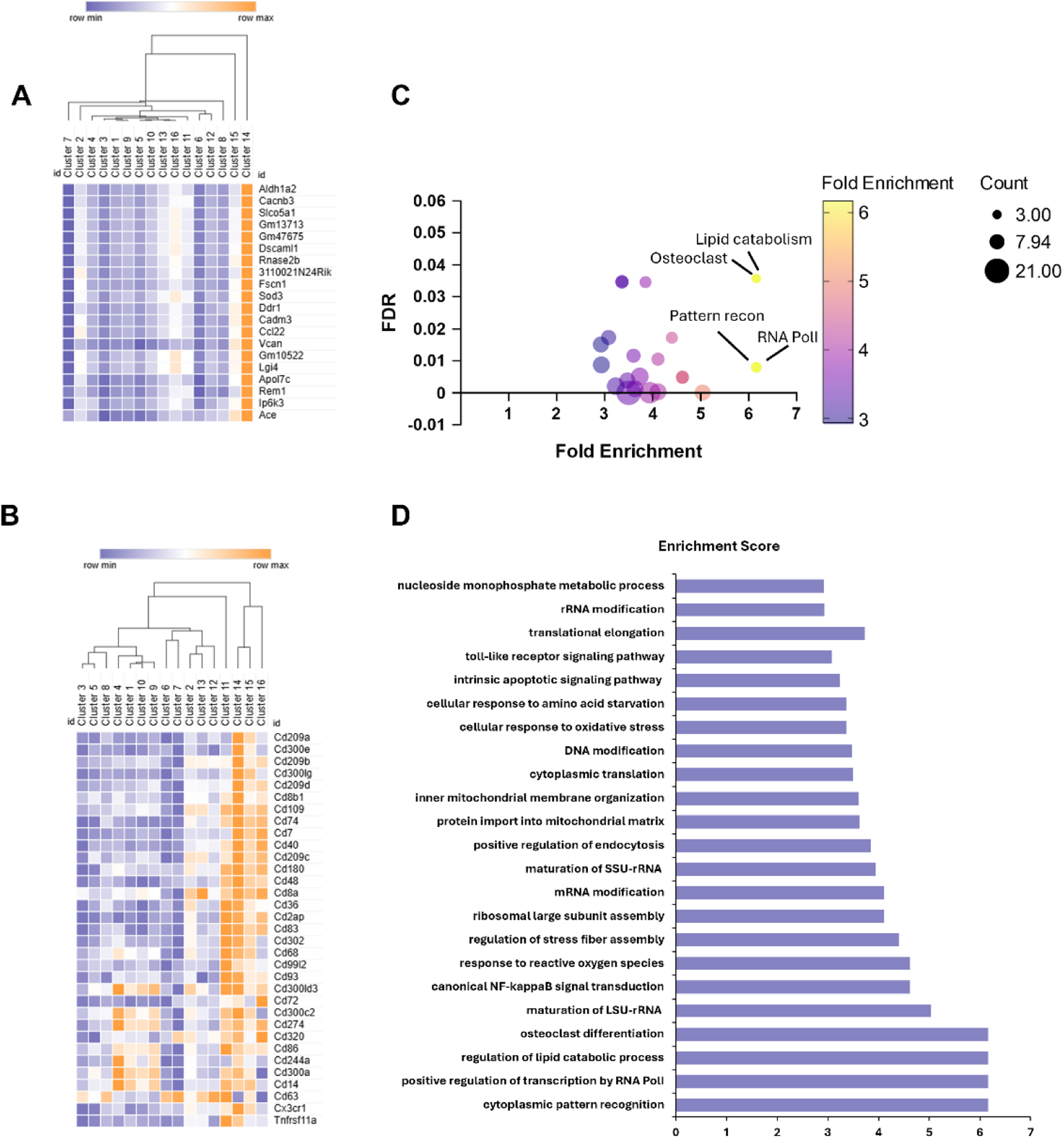
Cluster 14 Bioinformatic Analysis. (A) Heatmap of the top 20 differentially expressed genes within cluster 14 across all clusters. (B) Heatmap of surface markers defining cluster 14 across all clusters. (C) Bubble plot of enrichment, gene counts, and the false discovery rate for each functional annotation term. (D) GO biological process SLIM analyses were performed using differentially expressed genes within cluster 14. Shown are enrichment scores for each functional annotation.

We next sought to determine if our subsets of Cd74+ bone marrow-resident cells were also present under physiological conditions. We therefore profiled CD11B^+^ cells from 8-week-old male and female mice (Figure 6). Within males, we noted enriched expression of *Cd74* by cluster 9 (Figure 6A, B). These clusters were likewise marked by expression of *Tnfrsf11a* and *Acp5* (Figure 6A, B). When examining female CD11B^+^ cells, we likewise noted enriched expression of *Cd74* within clusters 3 and 11 (Figure 6C, D). Cluster 3 was further distinguished by differential expression of *CD163* and *CD36*, whereas *Tnfrsf11a* and *Acp5* marked cells within cluster 11 (Figure 6C, D). These data suggest that Cd74 may mark osteoclast progenitors within bone marrow of 8-week-old male and female mice under physiological conditions.

**Figure 6.**
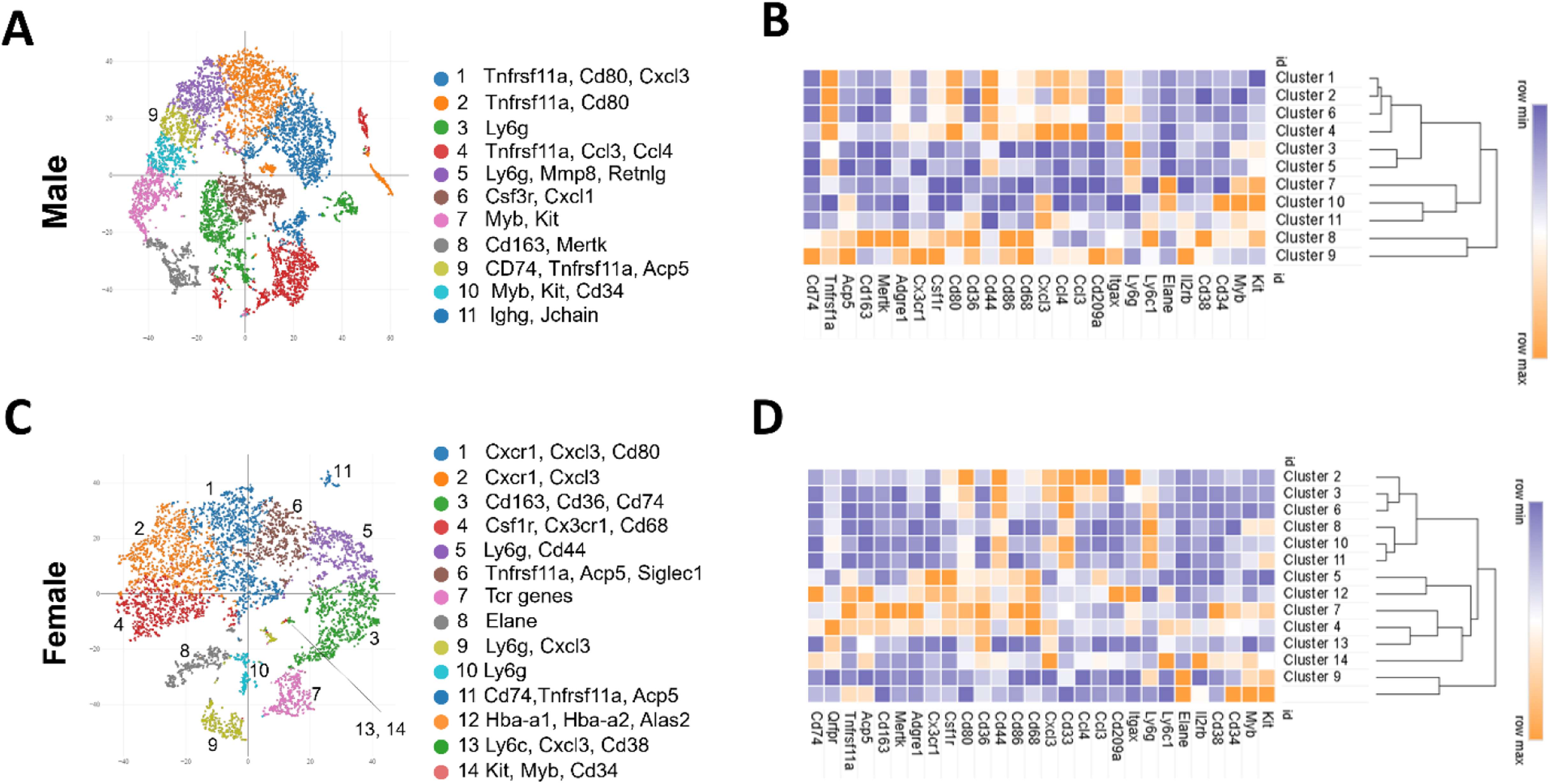
Single cell RNA-Seq of CD11B^+^ cells from male and female bone marrow. (A-D) CD11B^+^ cells from the bone marrow male and female mice were collected and single cell RNA-Seq was performed. (A) tSNE representation of data obtained from males. (B) Heatmap and hierarchical clustering of differentially expressed genes from clusters yielded from male data. (C) tSNE representation of data obtained from females. (D) Heatmap and hierarchical clustering of differentially expressed genes from clusters yielded from female data.

*Cd74* was the most variable feature within our scRNA-Seq data set and numerous *Cd74*-expressing clusters were differentially represented (Figure 3A, C, D). We therefore isolated *Cd74*-expressing cells within our dataset (Figure 7A). Clusters 2, 12, 15, and 16 demonstrated differential numbers of CD74-expressing cells, with the number of CD74-expressing cells lower in mice transplanted with female bone marrow (Figure 7B). We then collected nucleated bone marrow cells from 6-9-week-old C57Bl6/J mice and sorted cells into CD74^-^ and CD74^+^ fractions. While there were no differences between males and females within these fractions (Figure 7C, D), the percentage of CD74^+^ cells that were also CD11B^+^ was lower in females as compared to males (Figure 7E).

**Figure 7.**
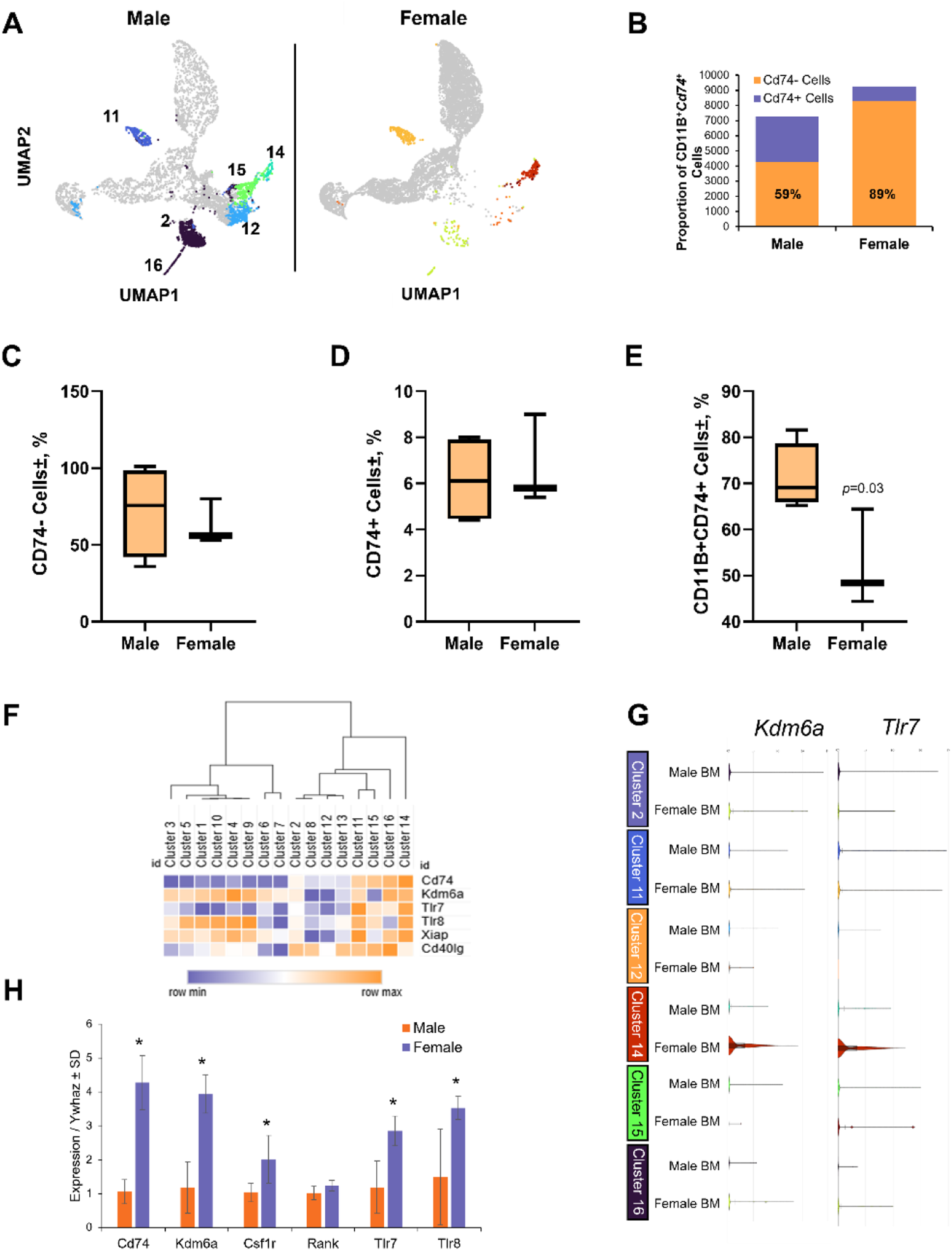
CD74 identifies osteoclast precursor-enriched populations associated with sex-specific differences. (A-C) Male recipient mice were lethally irradiated transplanted with T cell-depleted bone marrow (BM) from age matched males or age-matched female donor mice. Recipient mice were aged to 24-weeks for subsequent analyses. CD11B^+^ cells were collected from mice and scRNA-Seq was performed. Resulting data were integrated and comparative analyses were performed. (A) UMAP 2D representation of *Cd74*-expressing cells within CD11B^+^ clusters derived from recipient mice. (B) Proportion of *Cd74*-expressing cells as compared to total cell numbers. (C-D) Nucleated bone marrow cells were collected from 8-week-old male (n=4) or female (n=3) C57Bl6/J mice and CD11B^+^CD74^+^ cells were collected. Flow cytometry was performed to evaluate the (C) % Cd74^-^ cells, (D) % CD74^+^ cells, and (E) % CD11B^+^CD74^+^ cells. *p* values are as shown. (F) Heatmap of *Cd74* and X-linked genes across all clusters. (G) Expression of *KdmCa* and *Tlr7* by male and female cells within *Cd74*-expressing cells. (H) Real-time qPCR.

Our data suggest that cell-intrinsic differences in osteoclast progenitors may drive enhanced bone mass observed with transplantation of female bone marrow. Cell-intrinsic differences between males and females are in part driven by genes that escape X chromosome inactivation (XCI) in females, with the level of expression from the inactive X chromosome in women being a strong predictor of male-female differences (18, 19). The pattern and percent of XCI escape is variable between individuals and unique to tissue type (20). We therefore examined genes known to escape XCI that also regulate inflammation and/or osteoclastogenesis within *Cd74*-expressing clusters. X-linked genes *KdmCa*, *Tlr7/8*, *Xiap*, and *Cd40lg* were enriched within clusters 11 and 14, but showed variable expression within clusters 2, 15, and 16 (Figure 7F). *KdmCa* and *Tlr7* were specifically increased within female CD11B^+^*Cd74*^+^ cells within cluster 14 (Figure 7G). We then isolated CD11B+CD74+ cells from male and female mice via MACS. Expression of *Cd74*, *KdmCa*, *Csf1r*, and *Tlr7/8* was enriched in female cells (Figure 7H).

Our single cell sequencing data suggests that CD11/*Cd74* marks subsets of osteoclast progenitors (Figure 3, RANK+ cells). CD74 is also highly expressed by hematopoietic stem cells (21). We therefore determined the osteoclastogenic potential of CD11B^+^CD74^+^ cells (Figure 8A, B). Nucleated bone marrow cells were collected from 6-9-week-old male and female mice and MACS sorted into CD11B^+^CD74^-^ and CD11B^+^CD74^+^ fractions. Cells were cultured in the presence of RANKL and M-CSF and TRAP stained on day 4. CD74 is known to limit osteoclastogenesis, and we observed a trend toward fewer osteoclasts when comparing CD11B^+^CD74^-^ cultures to CD11B^+^CD74^+^ cultures from either males or females. We did not observe a difference between the number of CD11B^+^CD74^-^TRAP^+^ cells isolated from males or females. In contrast, fewer CD11B^+^CD74^+^TRAP^+^ multinucleated cells were observed in cultures derived from females as compared to males.

**Figure 8.**
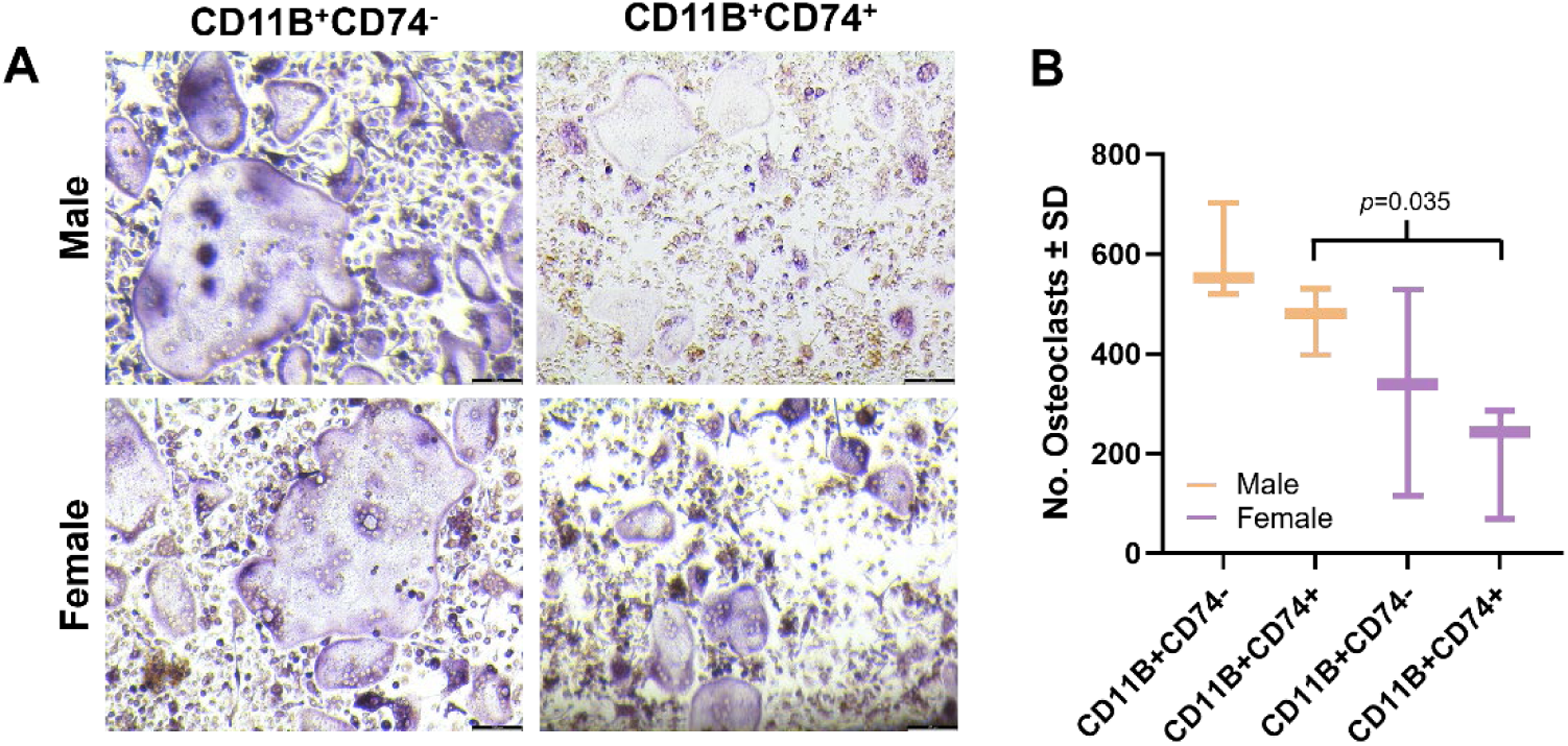
Assessment of osteoclastogenic potential in CD11b⁺CD74⁺ and CD11b⁺CD74⁻ progenitor cells from male and female mice. CD74 marks MACS-sorted cells were cultured in osteoclastogenic conditions, then TRAP stained (A) and enumerated (B) on day 4 (n=3). *p* values are as shown.

## Discussion

The goal of our study was to determine if cell intrinsic factors of bone marrow cells affected bone remodeling and bone mass. Clinical data and pre-clinical models demonstrate that females have lower vertebral bone mass (22). Our study demonstrates that transplantation of female cells enhanced vertebral body bone mass and was associated with changes in the cellular distribution of CD11B^+^ cells within the bone marrow. We also show microarchitectural changes following transplantation of female bone marrow cells into male hosts. Moreover, we show that *Cd74* marks osteoclast progenitors and that male CD11B^+^CD74^+^ cells have higher osteoclastic potential than female cells.

Despite the fact that bone mass is lower in females, cell-intrinsic properties of female transplanted bone marrow cells could enhance bone mass within a male host following lethal irradiation. Epigenetic programming differs between male and female HSCs (23). This may alter how transplanted bone marrow cells interact with male host osteoblast lineage cells. For instance, lineage negative female cells showed enhanced viability in a model of lung fibrosis (24). Likewise, changes in hematopoietic stem and progenitor cells with age are evident between males and females (25–29). Some studies also demonstrate that HSC and progenitor cell numbers are regulated by estrogen signaling (27, 28); thus, transplantation of female cells into male hosts could alter the number and potential of osteoclast progenitors independent of host circulating hormones via cell-intrinsic programming.

Using the four core genotypes model, mice harboring ovaries with an XX chromosomal content exhibited enhanced bone mass as compared to those with XY (10), demonstrating that cell-intrinsic differences due to chromosomal content control bone mass. These data support the notion that X-chromosome inactivation escape may drive differences between males and females. Within this study, expression of *Tlr7* and *KdmCa* by a population of CD11B^+^*Cd74*^+^ cells correlated with our observed differences in osteoclastogenesis and bone mass.

Differences in osteoclastogenic potential of male and female bone marrow in conditions of normal physiology have been reported in the literature, but the results have been conflicting. In murine models, osteoclast number was elevated and in vitro differentiation was accelerated using 8-week-old females as compared to males (30). In another study using 12-week-old mice, female bone marrow macrophages showed higher adhesion and proliferation in response to CSF1, while male cells exhibited increased responsiveness to RANKL (31). Using human peripheral blood cells, increased osteoclast differentiation and activity were observed in cultures derived from males; however, levels were comparable when comparing cultures derived from post-menopausal females and males (32). Likewise, male and female human osteoclasts exhibited differential responses to estrogens and androgens (33). Other studies likewise demonstrate differences in osteoclast differentiation and/or function between males and females (34–36). In our study, we found that numbers of CD11B^+^CD74^+^ cells were diminished in females. Moreover, we did not see a difference in osteoclastogenesis when comparing male and female CD11B^+^CD74^-^ cells, but male CD11B^+^CD74^+^ cells exhibited enhanced osteoclastogenesis when compared to females. This may potentially explain the enhancement in vertebral bone mass following transplantation with female cells, but further mechanistic studies are needed.

Another factor that could contribute to increased bone mass of male mice transplanted with female bone marrow cells is increased responses of the female osteoclast progenitors to estrogen signaling. Female monocyte/macrophage cells demonstrate higher estrogen receptor α and β levels and respond more strongly to estradiol (5, 37). Prior reports likewise demonstrate that estradiol reduces osteoclast numbers (38–40); thus, local amplification of estradiol signaling by transplanted female cells without a raise in serum estrogen in male hosts could contribute to our observed phenotype.

In our study we identified CD74 as a surface marker for osteoclast progenitors. Our scRNA-Seq dataset likewise supports that CD74^+^ cells are less abundant in male hosts transplanted with female cells. We validated this under conditions of normal physiology in 6-8-week-old mice. CD74 is an antigen-presenting molecule, but also functions as the receptor for macrophage inhibitory factor (MIF) (41–43). CD74 is expressed by immune cells, including cells within the monocyte/macrophage lineage and osteomorphs (43, 44). The CD74-MIF signaling axis reduces osteoclastogenesis by limiting NF-κB and c-Fos activation to reduce responsiveness to RANKL (45–49). Interestingly, male *Cd74* germline knockout mice demonstrate diminished trabecular bone mass that is not observed in females (47). We identified two *Cd74*-expressing clusters that are specifically enriched within male mice transplanted with female bone marrow. Osteoclastogenesis within this population may be increased due to CD74-dependent signaling. Diminished estrogen levels associated with menopause also correlated with reduced *Cd74* expression by tissue resident macrophages (50), but we were unable to show that estradiol treatment induced *Cd74* expression by in vitro cultured osteoclast progenitors. Future studies could be aimed at understanding how transcription induced by the CD74-MIF axis differs between males and females and if osteoclast progenitors marked by CD74 differ in abundance in conditions of altered bone mass or estradiol levels.

Our study has several limitations of note. First, while microarchitectural skeletal gains were predominant within the axial skeleton (L4 vertebral body), our scRNA-seq profiles were generated from femoral myeloid CD11B^+^ cells. We pooled samples from multiple mice for our scRNA-Seq experiment which limits the rigor of our analysis; however, our follow-up expression studies, flow cytometry, and in vitro osteoclastogenesis assays help to combat this limitation. Site-specific heterogeneity between axial and appendicular bone marrow niches is well recognized, which may account for why femoral metaphyses exhibited shifts in trabecular number without a net change in BV/TV. Future work will determine whether CD74^+^ progenitor subsets display identical transcriptomic signatures in vertebral marrow niche. We used mice in our study, but more work is needed to determine if CD11B/CD74 serve as markers for osteoclast progenitor cells both within mice and humans of both sexes. We noted diminished osteoclastogenesis of CD11B^+^CD74^+^ cells from either male or females; this observation could be due to inherent differences within these cells, but could also be explained by binding of the anti-CD74 antibody and downstream activation of CD74-dependent signaling. We also only evaluated young mice within this study, and it is possible that age and post-menopausal status would alter our findings. We also do not know if CD74^+^ serves as a surface marker to distinguish differences between males and females or if it has a functional role in defining different osteoclastogenic potential between sexes.

A critical consideration in interpreting our findings is the use of lethal irradiation prior to bone marrow transplantation, which induces substantial microvascular damage, acute trabecular bone loss, and widespread disruption of the recipient osteoblastic and stromal niches (51–53). Consequently, the elevated vertebral bone mass observed in male recipients reconstituted with female bone marrow may partly reflect sex-specific divergences in hematopoietic stem and progenitor cell radioprotection, homing efficiency, or lineage reconstitution kinetics during emergency hematopoiesis, rather than purely steady-state differences in baseline bone remodeling. Under conditions of post-radiation niche repair, rapid repopulation of the myeloid compartment could alter osteoclast precursor expansion and differentiation kinetics differently in female-derived donor cells compared to male-derived cells. Although our characterization of intact, non-irradiated 8-week-old mice confirms that females naturally harbor a smaller proportion of CD11B+CD74+ cells with reduced osteoclastogenic capacity, future studies using non-myeloablative conditioning or fate-mapping models will be critical to fully distinguish steady-state cell-intrinsic bone regulation from sex-specific responses to radiation-induced niche injury.

A notable observation in our study is the apparent divergence between our static bone histomorphometry results *in vivo* and our *in vitro* functional assays. While male hosts reconstituted with female bone marrow exhibited a 30% increase in vertebral bone volume, static histomorphometry at 24 weeks post-transplantation revealed no significant differences in osteoclast density (No. Oc/B.Pm) or osteoblast density (No. Ob/B.Pm). This suggests that the observed gain in bone mass may not stem from a sustained reduction in total osteoclast numbers at endpoint, but rather from functional alterations in individual osteoclast resorptive activity, cell fusion capacity, lifespan, or specific subsets of osteoclast progenitors. Alternatively, transient differences in osteoclast precursor differentiation and bone resorption kinetics may have occurred at earlier time points during post-irradiation hematopoietic reconstitution and skeletal accrual, settling into steady-state cell densities by 24 weeks. Because static parameters cannot capture dynamic bone remodeling kinetics over time, future studies incorporating fluorochrome double-labeling for dynamic histomorphometry (MAR and BFR/BS) as well as direct pit-resorption assays will be crucial to clarify whether female CD11B+CD74+ myeloid cells alter individual osteoclast resorptive efficiency or uncouple osteoclast-osteoblast signaling.

Overall, our study helps to define how cell intrinsic differences between male and female bone marrow cells help to control vertebral bone mass. We also identified CD74 as a surface marker for cells with osteoclastogenic potential. Our studies likewise suggest that increased numbers of CD11B^+^CD74^+^ cells correlate with reduced bone mass and that CD11B^+^CD74^+^ cells may dictate differences in osteoclast differentiation between males and females.

## ACKNOWLEDGEMENTS

The authors would like to thank the support from the Comparative Pathology Shared Resource, the Minnesota Dental Research Center for Biomaterials and Biomechanics, the University of Minnesota Genomics Core, and the University of Minnesota Flow Cytometry Resources (UFCR). This work was made possible by research grants from the National Institute for Musculoskeletal, Arthritis, and Skin Diseases (R21AR084530 to EWB), a seed grant to EWB from the UMN BIRCWH award (HD055887), and the University of Minnesota Board of Regents. These contents are solely the responsibility of the authors and do not necessarily represent the official views of the funding organizations. The study was conducted according to the guidelines of the University of Minnesota Institutional Animal Care and Use Committee (#2007-38247A approved on 08 August 2020). All data are contained within this manuscript and are available from the corresponding author upon reasonable request. Sequencing data from mice are deposited into the Gene Expression Omnibus (GEO accession numbers).

## AUTHOR CONTRIBUTIONS

JK and IYK contributed to investigation, writing of the original draft, editing, formal analysis and data curation. EKV, JCM, EP, SK, NR and MJS performed investigation, review and editing of the manuscript, formal analysis and data curation. KCM contributed to funding acquisition, reviewing and editing of the manuscript, and supervision. EWB contributed conceptualization, investigation, funding acquisition, writing of the original draft, review and editing, formal analysis and data curation.

---

**Supplemental Figure 1.**
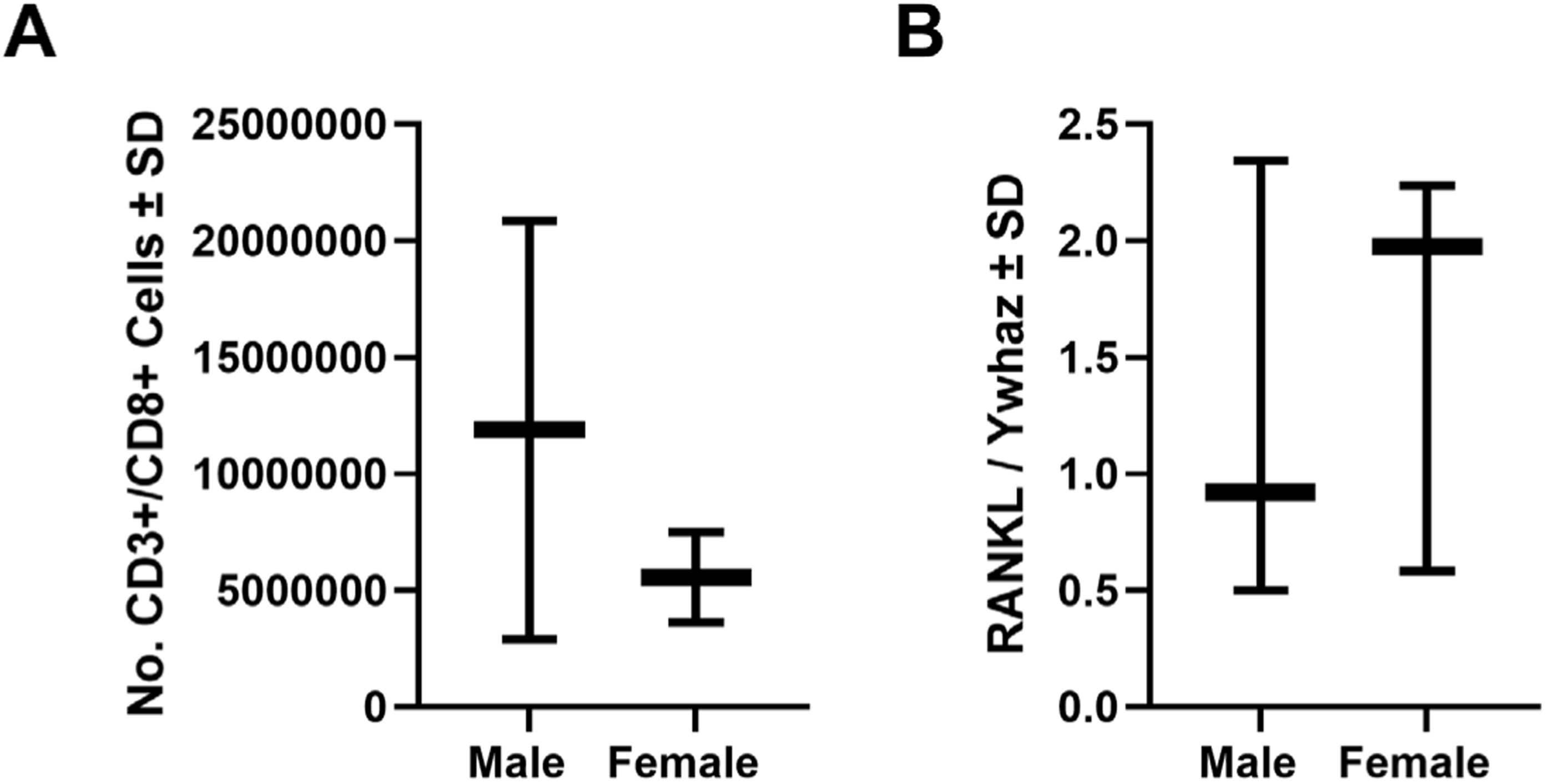
Male and Female CD3+CD8+ cells do not differentially express RANKL. (A) Numbers of CD3+CD8+ cells isolated from the bone marrow of male and female mice (n=3 per group). (B) Expression of RANKL by CD3+CD8+ cells isolated from males and females (n=3 per group).

**Supplemental Figure 2.**
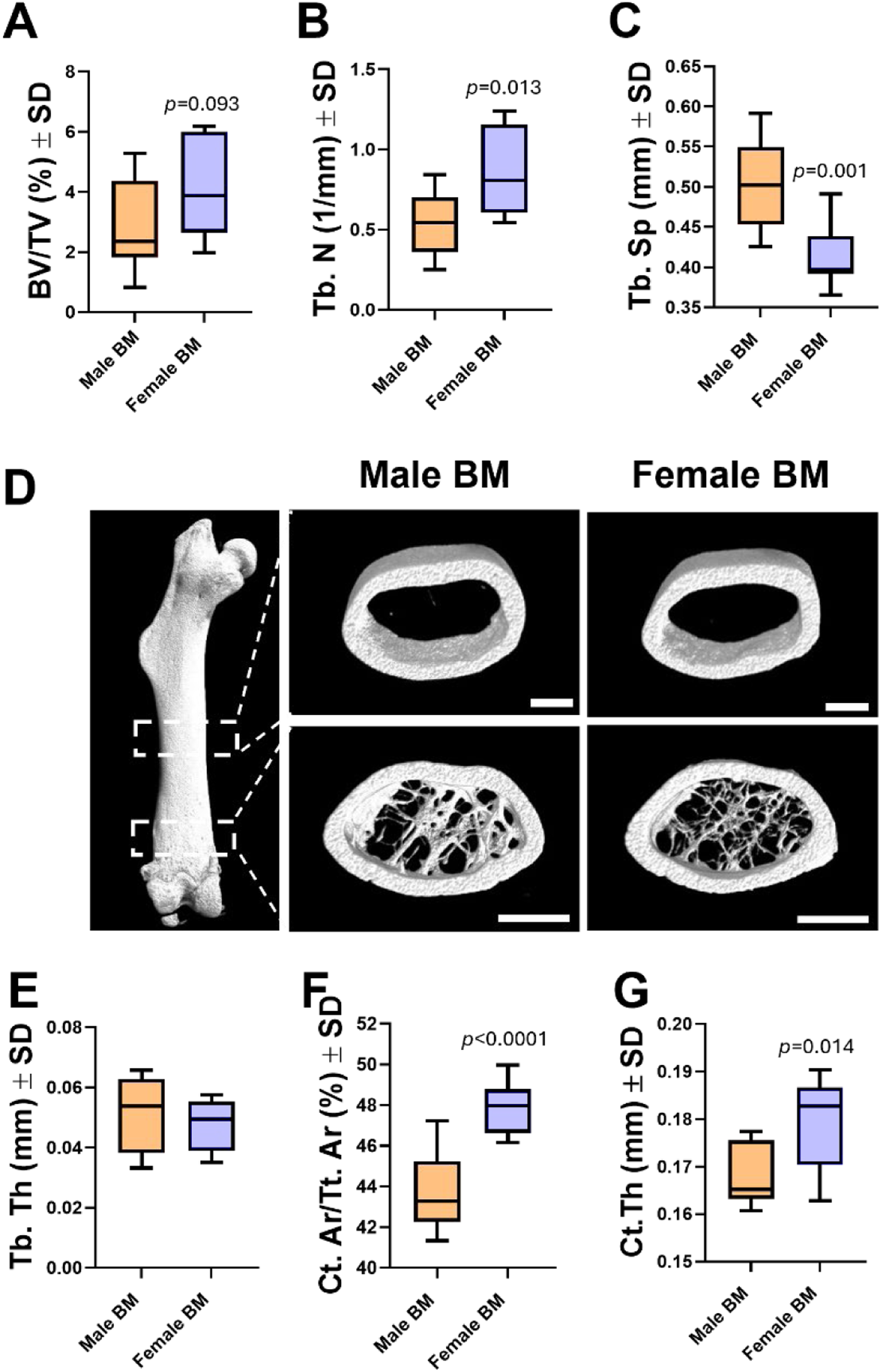
Femoral bone mass of male-female chimeric mice. Male-female chimeric mice display changes to femoral bone microarchitecture. Micro-CT analysis of the distal femur (A-C) or midshaft (E-G) was performed. (A) Bone volume per total volume (BV/TV), (B) Trabecular number (Tb. N), (C) Trabecular spacing (Tb. Sp), (D) reconstructions, (E) Trabecular thickness (Tb. Th), (F) Cortical area per total area (Ct.Ar/ Tt.Ar) and (G) cortical thickness (Ct. Th) are shown. *p* values under 0.05 are reported.

**Table 1.**
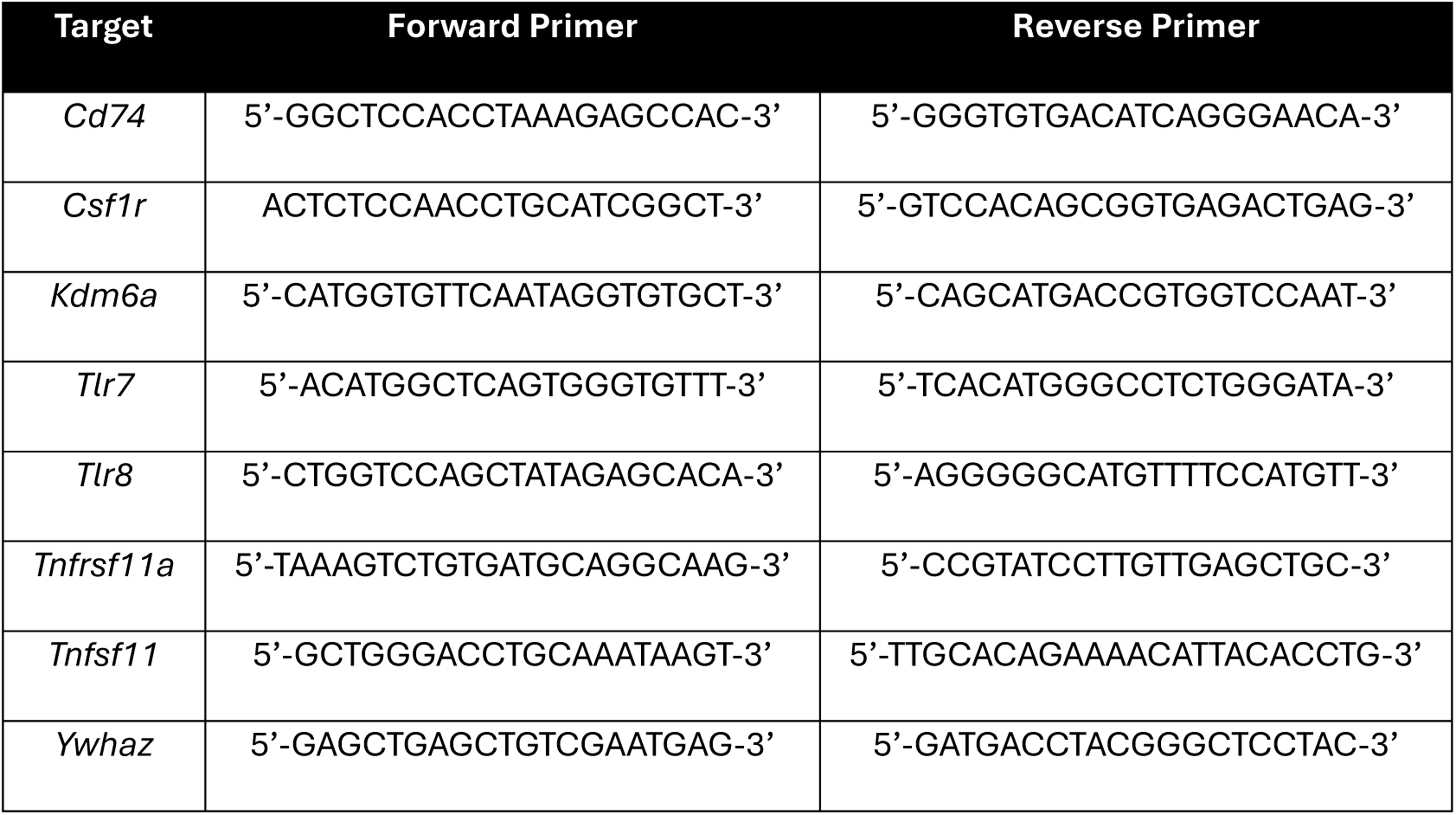
Primer Sequences.

